# Sensitivity to ATR–CHK1 pathway inhibition in AML/MDS is enhanced by *SRSF2* mutations and reduced by RUNX1 loss

**DOI:** 10.1101/2025.10.06.680457

**Authors:** Samuli Eldfors, Sumit Rai, Vineet Sharma, Tareq Hossan, Claudia Cabrera Pastrana, Amy Bertino, Angelique Gilbert, Kimmo Porkka, Matthew Walter, Timothy A. Graubert

## Abstract

*SRSF2* mutations occur in up to 25% of acute myeloid leukemia (AML) and 17% of myelodysplastic syndrome (MDS) cases and are associated with poor prognosis, yet no mutation-directed therapy exists.

Here, we aimed to identify therapeutically targetable vulnerabilities in MDS/AML with *SRSF2* mutations. Ex vivo drug-sensitivity testing of bone marrow cells from AML patients and healthy donors showed that *SRSF2*-mutant cells are sensitive to inhibitors of CHK1, and WEE1 DNA damage response (DDR) kinases.

To test causality, we engineered isogenic K562 cell line clones expressing *SRSF2*^P95H/L/R^ mutations. RNA sequencing confirmed splicing aberrations characteristic of MDS/AML in these clones. We found that *SRSF2*^P95H/L/R^ sensitize leukemia cells to ATR–CHK1–WEE1 inhibition. Bone marrow progenitors from *Srsf2*^P95H^ and *U2AF1*^S34F^ knock-in mice showed heightened sensitivity to CHK1 inhibition, corroborating the human data.

In contrast, *RUNX1* mutations were linked to resistance against CHK1 and WEE1 inhibition in *SRSF2*-mutant AML samples. *Runx1* loss also caused resistance to CHK1 inhibitors in knock-in mouse progenitors harboring *Srsf2*^P95H^ or *U2AF1*^S34F^, indicating that *RUNX1* loss is a mechanism of resistance.

In conclusion, *SRSF2* and *U2AF1* mutations are biomarkers of sensitivity to ATR–CHK1 pathway inhibitors, while *RUNX1* mutations cause resistance. These biomarkers can support patient stratification in MDS/AML.

## INTRODUCTION

Mutations in the *SRSF2* gene occur in 5–25% of patients with acute myeloid leukemia (AML) [1–3], in 17% of patients with myelodysplastic syndromes (MDS) [4], and up to 50% of patients with chronic myelomonocytic leukemia (CMML) [5]. The majority of *SRSF2* mutations result in substitution of P95 with histidine (H), leucine (L), or arginine (R). These hotspot mutations shift SRSF2’s RNA-binding preference from purine-rich GGNG to CCNG motifs, disrupting exon recognition in a sequence-dependent manner and causing widespread splicing alterations across numerous target genes. The most prominent defect involves altered inclusion or skipping of cassette exons [6,7].

*SRSF2* mutations have also been shown to impair transcriptional pause release by disrupting recruitment of P-TEFb from the 7SK snRNP complex, leading to increased formation of R-loops— RNA:DNA hybrids associated with genome instability and replication interference [6]. Given that splicing factor mutations such as *U2AF1*^S34F^ induce R-loop–dependent replication stress and create a dependency on the ATR-mediated DNA damage response pathway [8,9] it is plausible that *SRSF2* mutations might confer a similar vulnerability. However, whether the transcriptional and genomic instability caused by *SRSF2* mutations renders cells sensitive to pharmacologic ATR inhibition has remained unresolved.

Almost all patients with myeloid neoplasms that have *SRSF2* mutations have at least one additional gene mutation [7]. In patients with MDS and AML, mutations in *SRSF2* frequently co-occur with mutations in genes such as *IDH2*, *RUNX1*, *TET2*, *STAG2*, and *ASXL1* [1, 4, 8]. In contrast, *SRSF2* mutations are mutually exclusive with mutations in other splicing factor genes (*SF3B1*, *U2AF1*, and *ZRSR2)* and with mutations in genes including *TP53*, *EZH2*, and *DNMT3A* [9]. *SRSF2* mutations occur early in the development of MDS and AML [1, 8] and are, therefore, clonal and remain stable during disease progression [10]. These properties make mutant *SRSF2* a biomarker of interest and its downstream effects potential targets for therapy. *SRSF2* mutations are associated with adverse outcomes in MDS [10–13] and AML [1]. In MDS, *SRSF2* mutations are associated with older age and an increased risk of AML transformation [13]. However, despite adverse treatment outcomes, no targeted drugs exist for cancers with *SRSF2* mutations.

To address this unmet need, we aimed to identify therapeutically targetable vulnerabilities in MDS/AML with *SRSF2* mutations. Leveraging ex vivo drug-sensitivity testing—an approach that has successfully identified pharmacologically targetable vulnerabilities in multiple hematologic malignancies [14–16]—we analyzed drug sensitivity profiles of 109 primary AML patient samples screened against a library of approved and investigational oncology drugs. To validate the hits in a controlled experimental setting, we developed and characterized a panel of isogenic K562 cell line clones that express *SRSF2*^P95H/L/R^ mutations from the endogenous locus.

## METHODS

### Ex vivo Drug Sensitivity Testing of Patient Samples

Bone marrow mononuclear cells from AML patients and healthy donors were seeded into 384-well plates pre-loaded with compound library (five-point dilution series) and cultured for 72 h in RPMI medium supplemented with HS-5– conditioned factors [17]. Viability was measured with CellTiter-Glo (CTG) 2.0 assay and converted to drug-sensitivity scores (DSS) [18]. The screen, first reported in Ref. [16], was expanded over several years as with additional compounds; because the number of hypotheses changed between library versions, screen p-values are left unadjusted and are exploratory. DNA-damage–response inhibitors consistently ranked in the top 5% of the latest library and were therefore prioritized for validation. All patient material was obtained with written informed consent under Helsinki University Hospital approval, in accordance with the Declaration of Helsinki.

### Mutation and Chromosomal Aberration Identification

Gene mutations were identified by whole exome sequencing of leukemic bone marrow and matched skin biopsies using exome library preparation kits and sequenced on a HiSeq 2500 instrument. Somatic mutations were identified as described [14]. Chromosomal aberrations were identified using clinical karyotype data from the Finnish Hematology Registry and Biobank, Hospital District of Helsinki, Finland.

### Generation of Isogenic Gene-Edited Cell Lines

*SRSF2*^P95H/L/R^ mutations were introduced into K562 cells using CRISPR/Cas9. The ssODN includes a synonymous PAM-disrupting substitution to prevent Cas9 re-cutting. Clonal lines were expanded and confirmed by next-generation sequencing. Isogenic *SRSF2*^WT^ clones were used as controls. Sequences of primers, guides, and templates are in Supplementary Table S1.

### Cell Viability Assays

K562 or KO52 cells were seeded in 96-well plates and incubated with drugs for 72 or 96 h. Viability was determined using the CellTiter 96® Aqueous One Solution Cell Proliferation Assay (MTS) or CTG, and the results were normalized for growth rate using the GR metrics R package [19].

### RNA Isolation and Sequencing

RNA was extracted from K562 cells using the RNeasy Plus Mini Kit. Libraries were prepared with the NEBNext Ultra II RNA Library Prep Kit and sequenced on the NovaSeq 6000. Mutant Allele Frequencies in RNA: Paired-end reads were aligned to the human reference genome (NCBI GRCh38) using the STAR aligner [20].

### Processing and Aligning RNA Sequence Reads for Differential Splicing Analysis

Fastq files were processed with Fastp [21]. Gene annotations for hg19 from merging of Ensembl 87 [22] and UCSC knownGene databases were generated and used in the study. Reads were aligned to hg19 using RSEM [23] followed by alignment of remaining reads with Tophat [24]. Bowtie (1.0.0) was the base aligner in both steps.

### Differential Splicing Analysis

Splicing changes were quantified using MISO v2.0 [25] with human (hg19) splice annotations downloaded from the website (https://miso.readthedocs.io/en/fastmiso/annotation.html). Percent Spliced In (PSI) was calculated, and differentially spliced events were defined by thresholding of isoform ratio difference at 10% and having a Bayes Factor of 5 using Wagenmaker’s Bayesian framework [26]. Exonic splicing enhancer motif enrichment was analyzed using Biopython [27] (Detailed methods in supplementary). Visualization of Splicing Events: Sashimi plots were generated with MISO v2.0 [26] using the merged alignment files.

### Mouse models and induction

Conditional *Srsf2*^P95H/+^ [28] and *U2AF1*^S34F/+^ [29] knock-in mice, each bred to Mx1-Cre on C57BL/6J background, together with doxycycline-inducible *U2AF1*^S34^ and *U2AF1*^WT^ transgenics [30], were studied. Cre recombination was triggered by intraperitoneal plpC (12 µg g⁻¹) on days 0, 2 and 4; Tet-ON transgene expression was induced with doxycycline chow (625 ppm) for 5 days. All animal work was approved by the Washington University IACUC and conformed to NIH guidelines.

### Mouse bone marrow cultures

c-Kit⁺ progenitors were magnetically enriched from harvested bone marrow and cultured in RPMI-1640 containing 20% FBS, antibiotics, and a standard cytokine mix (SCF, TPO, FLT3-L, IL-3). CRISPR/Cas9 *Runx1* depletion: Bone marrow c-Kit⁺ progenitors were electroporated with Cas9– sgRNA ribonucleoproteins targeting *Runx1* (sgRunx1, 5′-TACCTGGTTCTTCATGGCCG-3′) or a non-targeting guide (NTG, 5′-GCTTTCACGGAGGTTCGACG-3′). GFP-tagged Cas9 V3 (IDT) and chemically modified sgRNA were pre-assembled (1:2.5 molar) and delivered with a Neon device; GFP⁺ cells were sorted 24 h later. Editing efficiency was quantified by amplicon sequencing and analyzed with CRISPResso2.

### Lentiviral transduction of bone marrow cells

Third-generation lentivirus encoding sgRunx1 or a non-targeting guide was produced in HEK293T cells and concentrated with Lenti-X. Bone marrow c-Kit⁺ cells were spin-infected at MOI 20 (1200 ×g, 2 h, 30 °C) in the presence of 5 µg/ml polybrene and cultured for 24 h before RFP⁺ sorting.

### Immunoblotting

Cell lysates were prepared in radioimmunoprecipitation assay (RIPA) buffer with protease and phosphatase inhibitors, clarified, and protein content equalized bicinchoninic acid (BCA) assay. Samples were mixed with 4× lithium dodecyl sulfate (LDS) sample buffer and β-mercaptoethanol, heated 10 min at 95 °C, separated by SDS–PAGE, and transferred to polyvinylidene difluoride (PVDF) membranes. Membranes were blocked in 5% milk/TBS-T, probed with anti-RUNX1 and β-actin, followed by HRP-conjugated secondaries and ECL detection.

### Cell viability and ex vivo drug sensitivity testing of mouse cells

Mouse bone marrow cell viability was measured using CTG. Cells were plated at 10,000 cells per well in 96-well plates. Drug sensitivity was assessed by comparing viability to a DMSO control.

Detailed experimental procedures can be found in the Supplementary Methods section available in the Supplementary Information.

### Accession Numbers

The Gene Expression Omnibus (GEO) accession number for the RNA sequencing data reported in this study is GSE213002.

## RESULTS

We reanalyzed ex vivo drug-sensitivity and somatic-mutation datasets generated by the Individualized Systems Medicine consortium [16] on bone marrow mononuclear cell (BM-MNC) samples from 109 AML patients and healthy donors. Ten samples (9%) harbored *SRSF2* mutations; *U2AF1* (3%) and *SF3B1* (2%) were less frequent, so subsequent analyses focused on *SRSF2*-mutant AML. Clinical characteristics are in Table 1; *SRSF2*-mutant cases were, on average, older at sampling, with other features comparable to *SRSF2* wild-type cases. Common co-mutations included *IDH2*, *RUNX1*, *STAG2*, and *TET2* (Figure 1), consistent with prior reports [1, 31].

**Figure 1.**
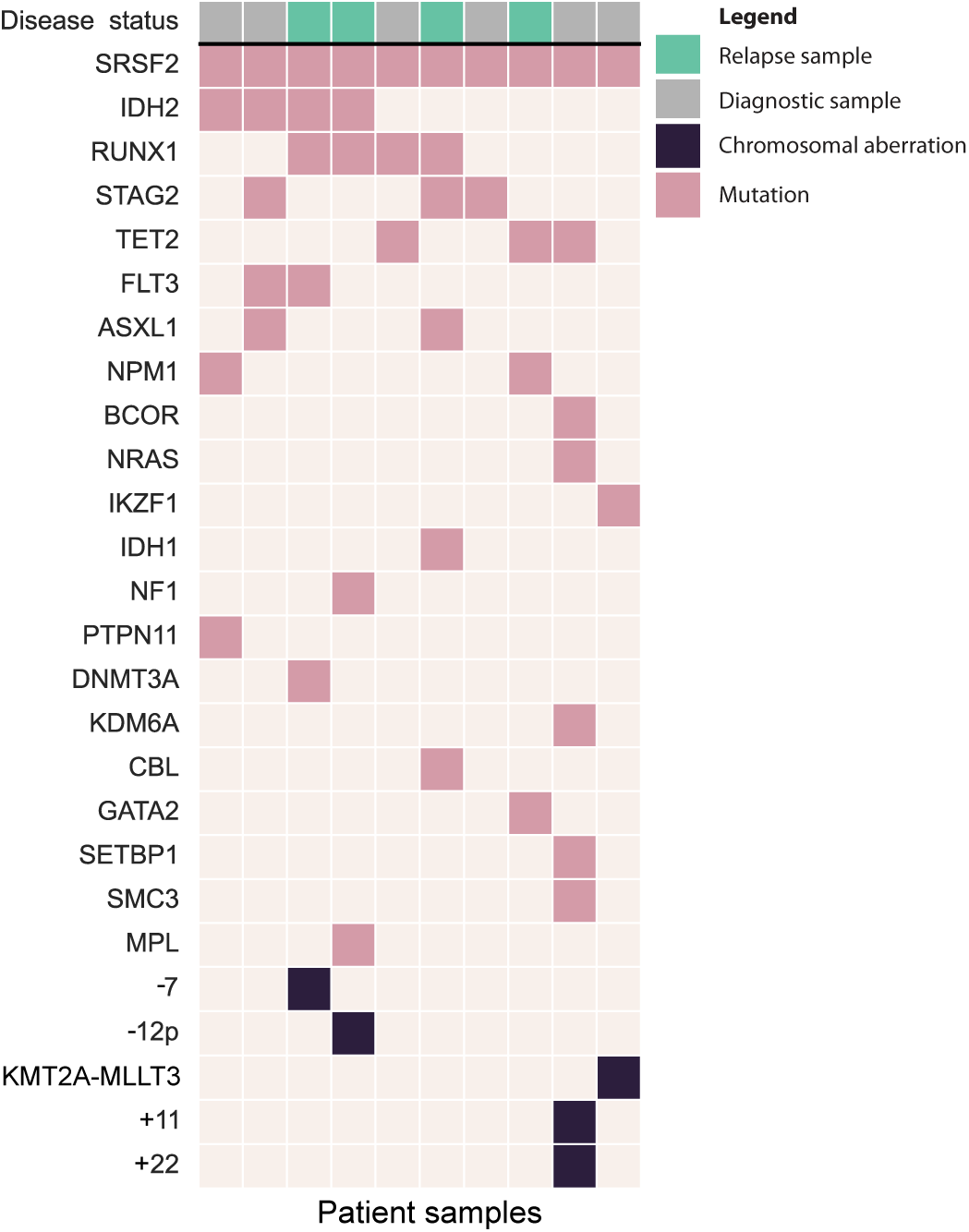
A Co-mutation plot showing gene mutations and chromosomal aberrations in bone marrow biopsies from 10 AML patients with *SRSF2* mutations. Chromosomal aberrations were detected through clinical karyotyping, while somatic mutations were identified via exome sequencing of leukemic bone marrow mononuclear cells and a matched skin biopsy.

**Table 1.**
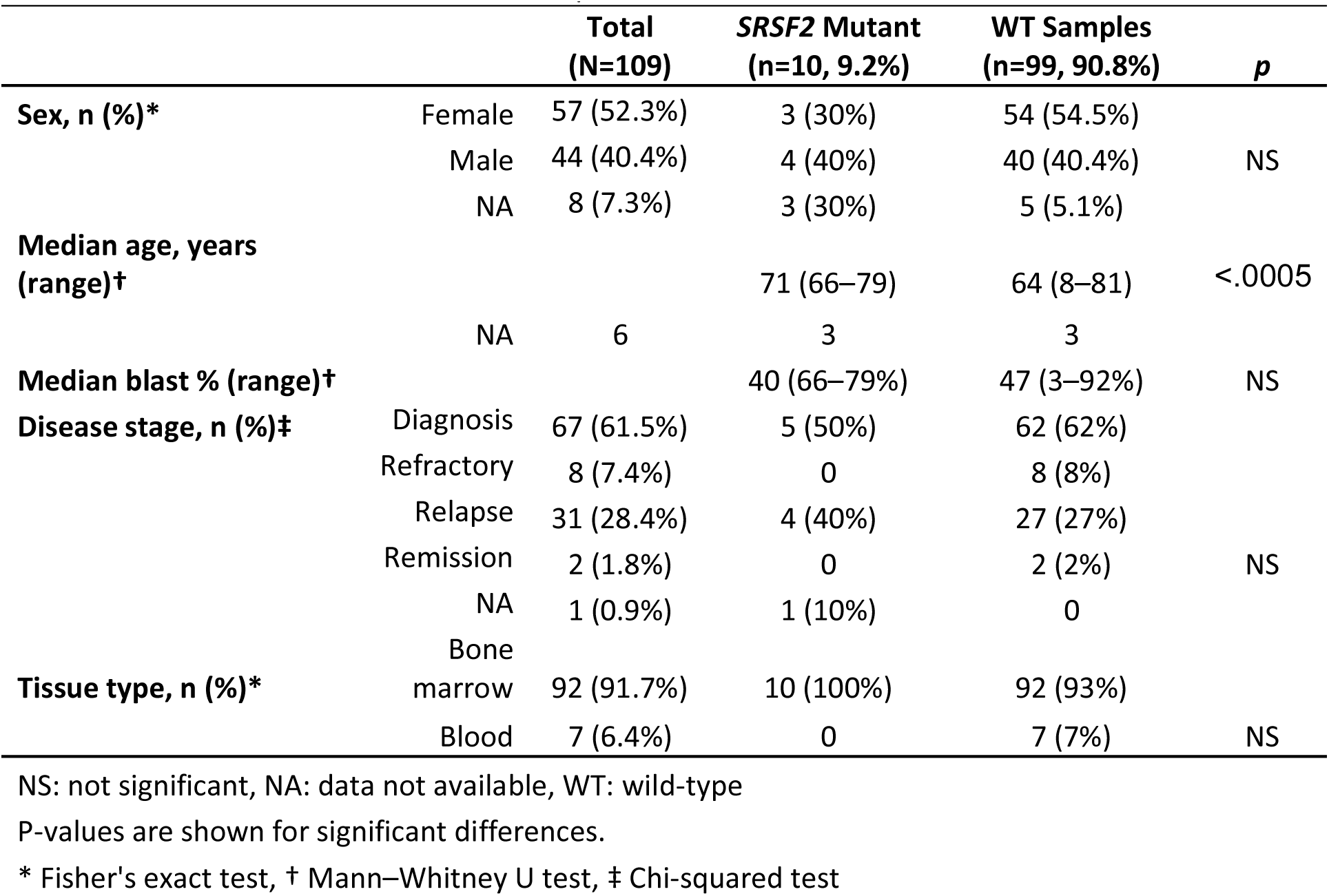
Clinical Characteristics of AML Sample Cohort.

### AML samples with *SRSF2* mutations exhibit sensitivity to ATR/CHK1/WEE1 inhibition, while *RUNX1* co-mutations are linked to resistance

Ex vivo drug-sensitivity data from primary AML BM-MNCs ranked CHK1 and WEE1 inhibitors among the top 5% compounds associated with *SRSF2* mutations. Given our previous finding that splicing factor mutations confer sensitivity to ATR inhibition [32], we examined the response of *SRSF2*-mutant AML samples to ATR inhibitors and its downstream effector kinases, CHK1 and WEE1.

Of the nine *SRSF2*-mutant AML samples, five showed striking hypersensitivity to the CHK1 inhibitor prexasertib (72-h IC_50_ 0.2–4.5 nM). The remaining four were resistant, showing an appreciable loss of cell viability only when exposed to the highest dose tested (1 µM; Figure 2A). The resistant AML samples were characterized by co-occurring clonal *RUNX1* mutations in three cases and a subclonal *SRSF2* mutation in one case (Figure 2A).

**Figure 2.**
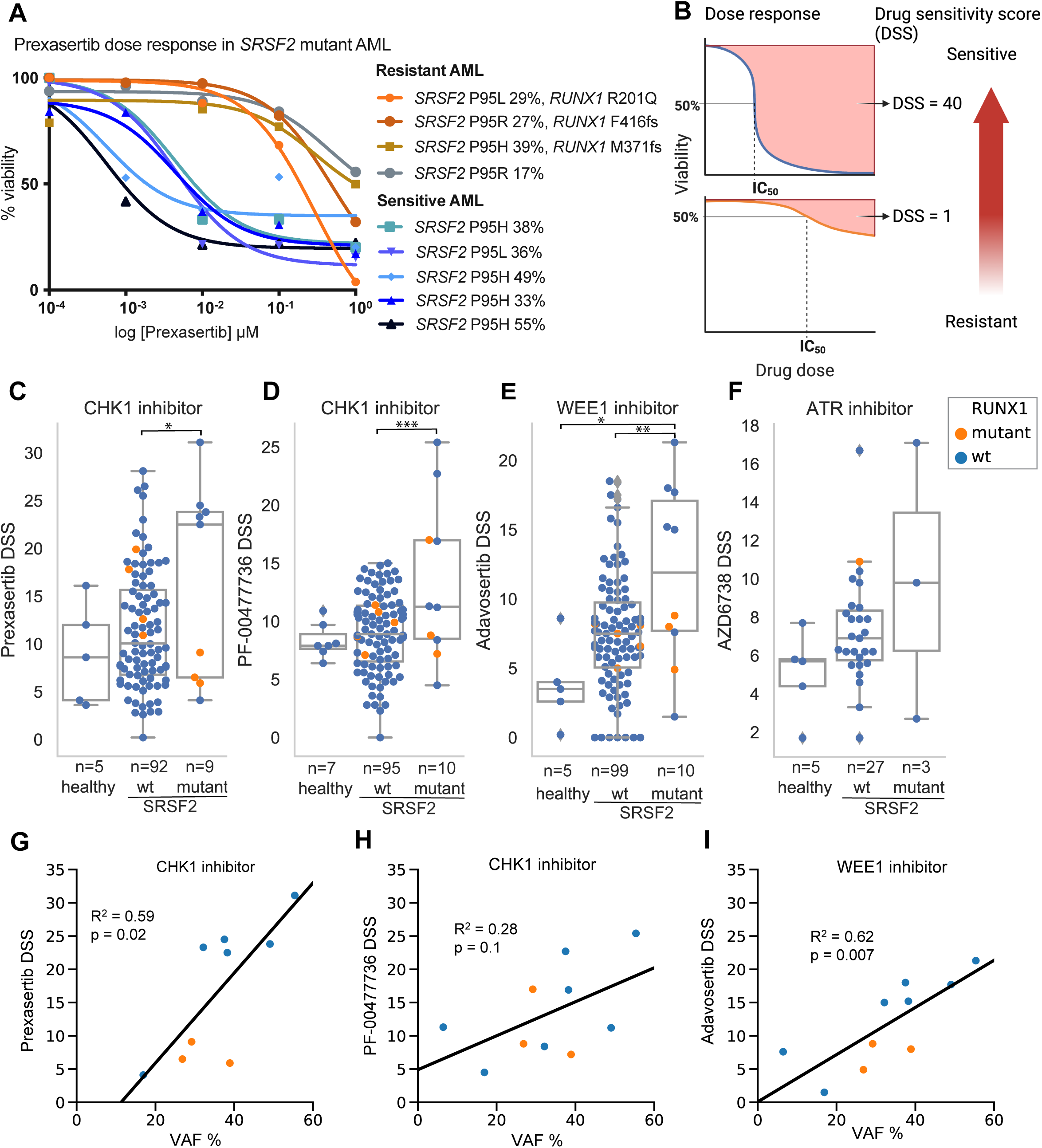
AML samples with *SRSF2* mutations are sensitive to inhibition of ATR/CHK1/WEE1 DNA damage response kinases. (A) Fresh bone-marrow mononuclear cells were treated for 72 h with prexasertib in five concentrations (0.001–10 µM); viability was measured by CellTiter-Glo and normalized to a DMSO (methods apply to A–I). (B) Schematic illustrating the relationship of dose-response curves and drug sensitivity scores (DSS). The area delineated by a dose-response curve is transformed into a DSS using an integral function described previously [18]; higher DSS indicates greater sensitivity. (C–F) DSS in healthy donors (healthy), *SRSF2*-wildtype AML (wt), and SRSF2-mutant AML (mutant) for CHK1 inhibitors prexasertib (C) and PF-00477736 (D), WEE1 inhibitor adavosertib (E), and ATR inhibitor ceralasertib (F). Points are individual patient samples; *RUNX1*-mutant samples are orange and *RUNX1*-WT blue. Horizontal lines indicate medians. P values by two-sided Mann–Whitney U test; P<0.05, *P<0.01, **P<0.001. (G–I): Linear regression of *SRSF2* variant-allele frequency (VAF, %) versus DSS for prexasertib (G), PF-00477736 (H), and adavosertib (I). Lines show least-squares fits.

Dose-response curves were converted into drug-sensitivity scores (DSS; Figure 2B) to facilitate comparison. Most *SRSF2*-wild-type AML samples showed only modest sensitivity to the CHK1 inhibitor prexasertib (median DSS = 11; range 0.2–28), which was markedly lower than in the *SRSF2*-mutant samples. In *SRSF2*-wild-type AML samples, the presence of a *RUNX1* mutation was not associated with resistance to prexasertib.

Across healthy donors, *SRSF2*-WT AML, and *SRSF2*-mutant AML, the mutant group showed higher DSS to CHK1 inhibitors, prexasertib (Figure 2C) and PF-00477736 (Figure 2D), and to the WEE1 inhibitor adavosertib (Figure 2E). Within the *SRSF2*-mutant group, adavosertib responses were bimodal: the most sensitive samples lacked *RUNX1* co-mutations, whereas less-sensitive samples harbored *RUNX1* co-mutations or had subclonal *SRSF2*, resembling the pattern observed with prexasertib. Sensitivity to the ATR inhibitor ceralasertib (AZD6738) showed a similar direction but was not tested statistically due to low number of *SRSF2*-mutant cases (Figure 2F).

We next evaluated whether the variant allele frequency (VAF) of *SRSF2* mutations was associated with response to CHK1 and WEE1 inhibitors using linear regression. *SRSF2* VAF correlated significantly with sensitivity to prexasertib and adavosertib in primary AML samples (Figures 2G, I). Although PF-00477736 showed a comparable trend, the association did not reach statistical significance (Figure 2H).

Taken together these data indicate that AML with clonal *SRSF2* mutations is vulnerable to inhibition of ATR, CHK1, and WEE1, while samples with co-occurring *RUNX1* mutations were resistant.

### Isogenic K562 cells carrying *SRSF2* mutations recapitulate splicing aberrations in AML and MDS

To study vulnerabilities caused by *SRSF2* mutations, we used CRISPR/Cas9 homology-directed repair to generate a panel of 11 isogenic K562 cell line clones expressing *SRSF2* P95H, P95L or P95R mutant allele from the endogenous *SRSF2* locus (Supplementary Table S2). The presence of mutant alleles was confirmed by amplicon sequencing (Figure 3A). K562 cells have three copies of *SRSF2*: one on chromosome 17 and two copies of a derivative chromosome der(17)t(9;17) [33]. This aneuploidy allowed us to generate K562 clones with a range of *SRSF2* mutant allele copy numbers. We utilized amplicon sequencing to infer the number of mutant alleles in each clone. We categorized VAFs into non-overlapping ranges: 1–33% corresponding to one mutant allele, 34–66% to two, and 67–100% to three. Expression of the mutant *SRSF2* transcript was detected in all mutant clones by RNA sequencing (Figure 3B).

**Figure 3.**
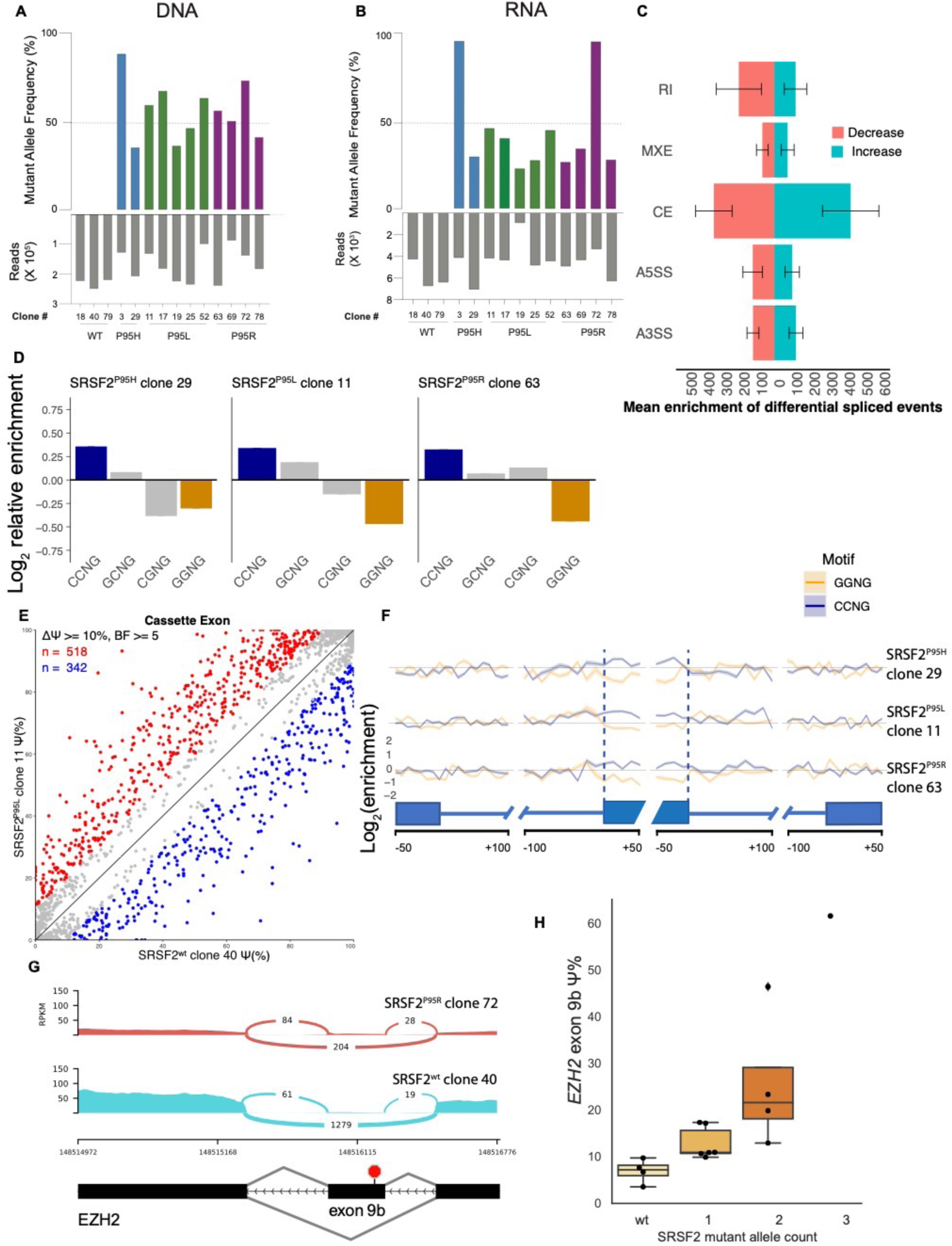
*SRSF2* mutations alter RNA splicing and exonic splicing enhancer preference in K562 cells. (A) *SRSF2*^P95H/L/R^ variant allele frequency (VAF) in isogenic K562 clones (3 wild-type and 11 mutant) measured by amplicon sequencing; gray bars indicate read depth. (B) Expression of *SRSF2* mutant alleles by whole transcriptome RNA-seq. (C) Differential splicing by event class in *SRSF2*-mutant K562 clones vs the isogenic WT control. Bars show the mean number of events per clone meeting the significance threshold; right = increased inclusion, left = decreased (colors as indicated). Categories: retained intron (RI), mutually exclusive exons (MXE), cassette exon (CE), alternative 5′ splice site (A5SS), alternative 3′ splice site (A3SS). Error bars, SD across 11 mutant clones. Significance thresholds as in panel E. (D) Mean enrichment of variants of the SSNG exonic splicing enhancer motif within cassette exons shows increased vs. decreased inclusion in isogenic K562 clones with *SRSF2*^P95H/L/R^ mutations compared to the wild-type control (’S’ represents G or C). The mean enrichment for a given motif was defined as the mean of the count of instances in all exons with increased inclusion divided by the count of instances in all exons with reduced inclusion. (E) Scatter plot of cassette exon inclusion comparing *SRSF2*^WT^ K562 clone 40 (x-axis) and mutant *SRSF2*^P95L^ clone 11 (y-axis). Percent-spliced-in (Ψ) indicates the proportion of cassette exon inclusion. Points represent individual cassette exons; red = higher inclusion in mutant vs WT, blue = lower, gray = not significant. Cassette exons with significantly different exon inclusion have |ΔΨ| ≥ 10 percentage points and a Bayes factor ≥ 5, as estimated by Wagenmakers’ framework [26]. Diagonal (y=x) indicates no difference. (F) Relative enrichment of CCNG and GGNG motifs in cassette exons with significantly increased vs decreased inclusion in three representative isogenic K562 clones with *SRSF2*^P95H/L/R^ across a metagene depicting differentially spliced exons. x-axis: nucleotide position relative to splice-site junctions; y-axis: log₂(increased/decreased) motif-count ratio (positive = enrichment among cassette exons with increased-inclusion). Lines are clone-specific; shaded bands denote 95% bootstrap confidence intervals. (G) Sashimi plot at the *EZH2* exon 9b locus. The *SRSF2^P95R^* clone (top; clone 72) shows greater exon 9b inclusion than the isogenic WT clone (bottom; clone 40). y-axis: normalized RNA-seq coverage (RPKM); x-axis: hg19 chr7 coordinates. Arc labels indicate counts of junction-spanning reads. (H) Ψ for *EZH2* exon 9b in isogenic K562 clones grouped by *SRSF2* mutant allele count [0 (WT), 1/3, 2/3, 3/3]. Boxes show the interquartile range (25^th^–75^th^ percentiles) with the center line median; points are individual clones.

To determine whether introducing *SRSF2*^P95H/L/R^ alleles into K562 cells recapitulates splicing aberrations observed in MDS/AML with *SRSF2* mutations, we performed high-depth RNA-sequencing on isogenic clones and an isogenic WT control (Supplementary Table S3) and quantified differential splicing. All classes of alternatively spliced events were altered in *SRSF2*-mutant clones, with the most prominent impact on cassette exon use (Figures 3C).

Given that *SRSF2* mutations alter binding affinities toward CCNG and GGNG exonic splicing enhancer (ESE) motifs, leading to altered exon inclusion [28, 34], we assessed whether these splicing changes are recapitulated in the isogenic K562 clones with *SRSF2* mutations. We identified differentially spliced exons in each clone and computed the enrichment of ESE motif variants within cassette exons that had increased vs decreased inclusion in mutant compared with a wild-type control clone. We found that all isogenic *SRSF2* mutant clones exhibited increased inclusion of cassette exons with CCNG motifs and reduced inclusion of those with GGNG motifs (Figures 3D– F and Supplementary Figures S1A,B).

*SRSF2* mutations have also been shown to cause increased inclusion of *EZH2* exon 9b into mature RNA. *EZH2* exon 9b is a conserved poison exon that contains a premature stop codon [28, 35]. We analyzed exon inclusion events in isogenic K562 clones and found higher exon 9b inclusion in *SRSF2* mutant clones compared to the wild-type controls (Figure 3G). *EZH2* exon 9b inclusion increased with the number of mutant *SRSF2* alleles (Figure 3H).

### *SRSF2*^P95H/L/R^ mutations sensitize K562 cells to ATR, CHK1 and WEE1 inhibitors

To determine whether *SRSF2* mutations are sufficient to sensitize leukemia cells to inhibition of ATR, CHK1, and WEE1, we tested the sensitivity of ten isogenic *SRSF2* mutant and four wild-type K562 clones to inhibitors of these kinases. These isogenic wild-type and mutant pairs were derived from the same electroporated pool, ensuring a consistent genomic background.

We determined the dose response of the clones to the ATR inhibitor elimusertib (BAY-1895344); CHK1 inhibitors, prexasertib (ACR-368, LY-2606368) and SRA-737; and the WEE1 inhibitor adavosertib (AZD-1775). These inhibitors were chosen for their demonstrated activity in short-term *in vitro* viability assays, good target specificity, and the fact that they have been studied in clinical trials [36, 37]. Cells were incubated with increasing concentrations of the inhibitors for 72 h, followed by the determination of cell viability using MTS assay. Since *SRSF2* mutations affect K562 growth, we used the GR algorithm to normalize viability for growth-rate differences [19, 38]. We found that *SRSF2*-mutant clones demonstrated a statistically significant increase in sensitivity to CHK1 inhibitors, prexasertib, and SRA-737 (Figures 4A–G). A similar trend of increased sensitivity was observed for WEE1 and ATR inhibitors.

**Figure 4.**
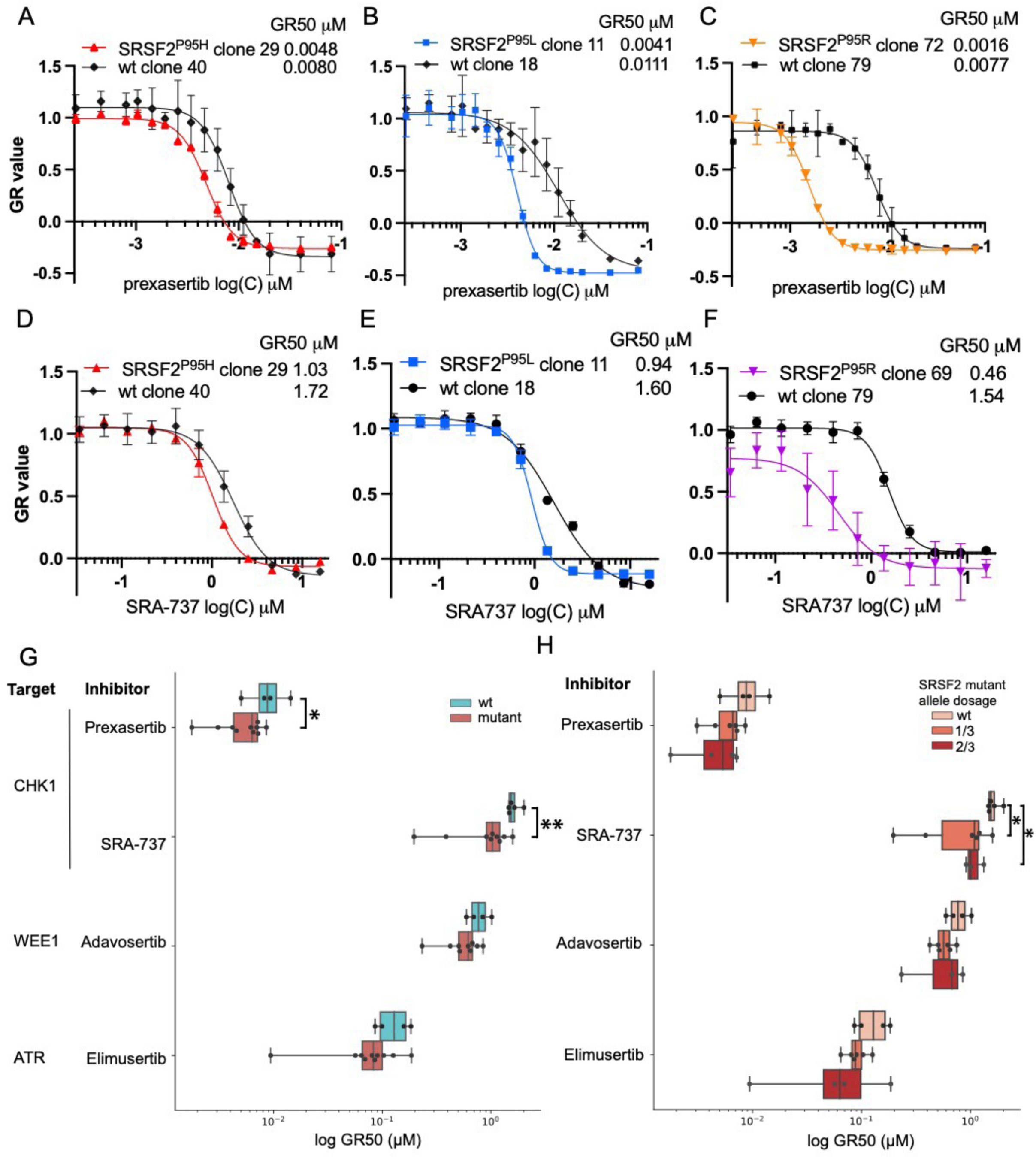
ATR/CHK1/WEE1 inhibitor sensitivity in isogenic K562 clones carrying *SRSF2*^P95H/L/R^ mutations. A–F: Cells were treated for 72 h; viability was measured by MTS and converted to growth-rate inhibition (GR). (A–C) Prexasertib dose-response of isogenic K562 cell line clones with *SRSF2* mutations P95H (A), P95L (B), P95R (C), and matched wild-type controls. (D–F) SRA-737 dose-response curves for the same clones and matched wild-type controls. (G) Summary across clones: log_10_(GR_50_, µM) for prexasertib, SRA-737, adavosertib, and elimusertib in mutant (n = 10) vs WT (n = 4) K562 clones. Points are drug GR_50_ values for individual clones. Horizontal bars show median IQR. *, P < 0.05; **, P<0.01 by unpaired two-sided Student’s t-test. (H) Log_10_(GR_50_, µM) by *SRSF2* mutant-allele copy number inferred from amplicon-sequencing VAF: 0 (WT), 1/3, 2/3.

Next, we sought to determine whether our findings could be replicated using an alternative assay. We performed a dose-response analysis with the CHK1 inhibitor prexasertib using the Incucyte Live-Cell Imaging system. *SRSF2*^P95L^ mutant cells and their corresponding wild-type controls were exposed to prexasertib for 96 h. Post-incubation, cell counts, and dose-response curves were plotted to derive IC_50_ values. Our analysis revealed that the IC_50_ for *SRSF2* mutant cells was notably lower than for the wild-type controls, reconfirming the increased sensitivity of *SRSF2* mutant cells to prexasertib (Supplementary Figure S2).

K562 cells contain three copies of the *SRSF2* gene, and the isogenic clones have 1, 2, or 3 mutant allele copies. We hypothesized that a higher *SRSF2* mutant allele number causes an increased dependence on the ATR pathway. Therefore, we asked whether a higher number of *SRSF2* mutant alleles leads to increased sensitivity to ATR/CHK1/WEE1 inhibition. We found that clones carrying two mutant *SRSF2* alleles had increased sensitivity compared to a single mutant allele, consistent with our prediction (Figure 4H). A single clone with three *SRSF2* mutant alleles had a profoundly reduced growth rate, and the dose response could not be evaluated.

To determine whether the specific missense substitutions influence drug sensitivity, we compared the sensitivities of clones grouped by *SRSF2*^P95H/L/R^ substitution. There was no significant difference in sensitivity to ATR/CHK1/WEE1 inhibition between clones carrying *SRSF2*^P95H/L/R^ mutant alleles (Supplementary Figure S3).

Together, these data demonstrate that *SRSF2* mutations sensitize K562 cells to ATR, CHK1, and WEE1 inhibitors, with the largest effect for CHK1 inhibitors.

### Runx1/RUNX1 loss confers CHK1-inhibitor resistance in splicing-factor–mutant mouse hematopoietic progenitors and human leukemia cells

Because AML samples from patients with concurrent *SRSF2* and *RUNX1* mutations were resistant to CHK1 inhibition (Figure 2A), we tested whether *Runx1* depletion confers CHK1-inhibitor resistance in splicing-factor–mutant mouse hematopoietic progenitors.

To test this, we used knock-in (KI) mouse models carrying either *Srsf2*^P95H^ or *U2AF1*^S34F^, each with matching wild-type controls. Bone marrow c-Kit⁺ progenitor cells were isolated and nucleofected with Cas9-GFP ribonucleoprotein (RNP) complexes assembled with either non-targeting (NTG) or *Runx1*-targeting sgRNA. Sorted GFP⁺ cells were exposed ex vivo to the CHK1 inhibitor SRA-737 for 96 h across a concentration range to generate dose-response curves. Editing and protein depletion was confirmed by amplicon sequencing and western blotting (Supplementary Figures S4A,B and S6A,B).

Inducing *Srsf2*^P95H^ increased SRA-737 sensitivity, lowering IC₅₀ ∼3.2-fold relative to *Srsf2*^WT^ (0.19 vs 0.63 µM; P=0.0014). *Runx1* depletion made the *Srsf2*^P95H^ cells more resistant, increasing IC_50_ to ∼0.52 µM (∼2.9-fold vs NTG; (p = 0.009). One-way ANOVA with Tukey correction (Figure 5A and Supplementary Figure S4B).

**Figure 5.**
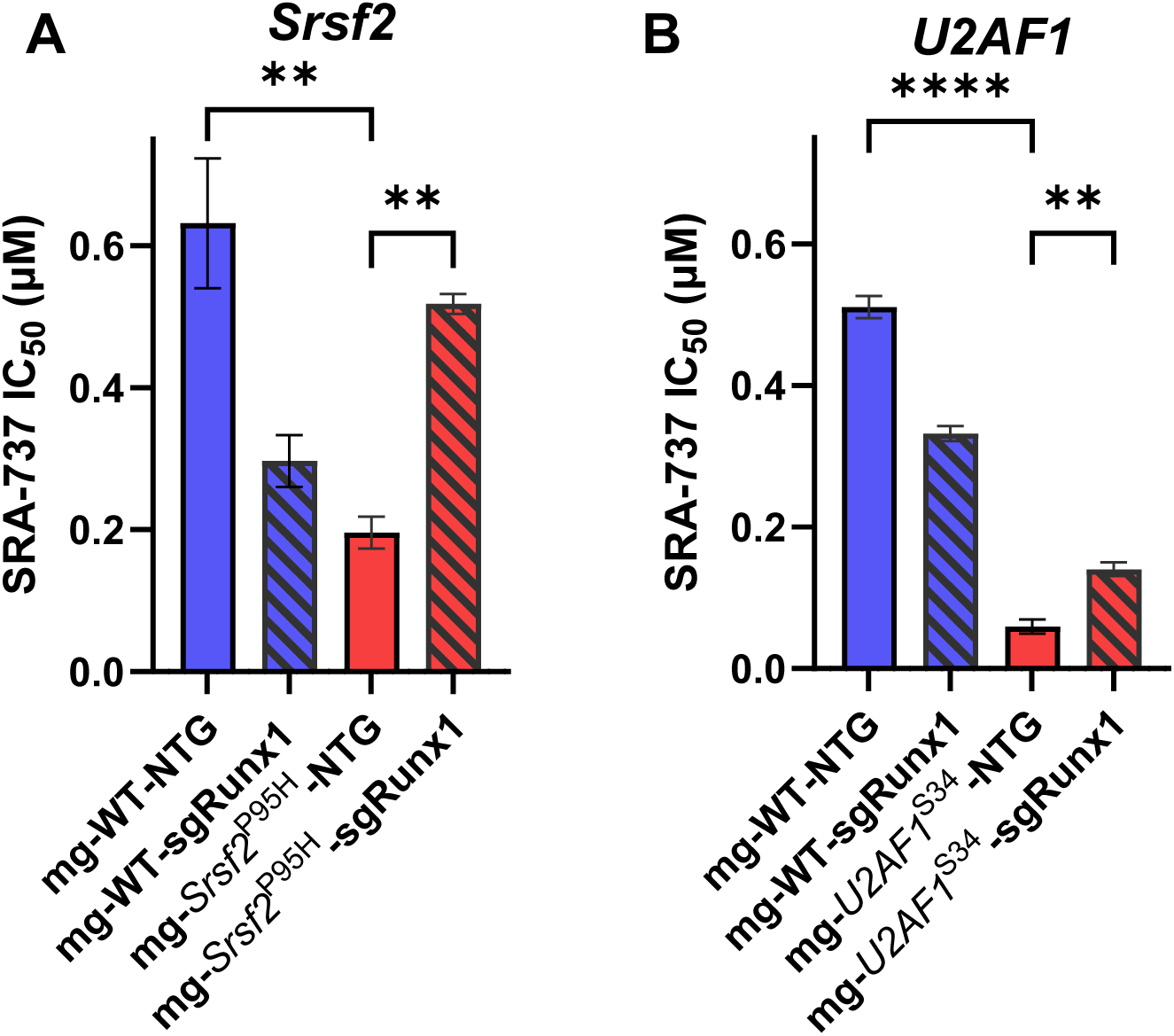
*Runx1* confers resistance to SRA-737 in splice-factor–mutant c-Kit⁺ progenitors. Primary mouse bone marrow c-Kit⁺ hematopoietic progenitor cells were treated with SRA-737 for 96 h and viability was measured with CellTiter-Glo 2.0. Bars show mean IC_50_ (µM) ± SD from n = 3 technical replicates per genotype. Color/pattern mapping: WT-NTG (solid blue), WT-sgRunx1 (hatched blue), mutant-NTG (solid red), mutant-sgRunx1 (hatched red). Statistics were performed on replicate IC_50_ values by one-way ANOVA with Tukey’s multiple comparisons; **, P<0.01; ****, P<0.0001). Lower IC_50_ indicates greater sensitivity. Panels: (A) *Srsf2*^P95H^, (B) *U2AF1*^S34F^.

In *U2AF1*^S34F^ cells, NTG controls were 8.5-fold more sensitive to SRA-737 than *U2AF1*^WT^ (IC_50_: 0.060 µM vs. 0.51 µM, p<0.0001)*. Runx1* depletion made the cells more resistant, increasing the IC_50_ of *U2AF1*^S34F^-sgRunx1 cells 2.4-fold compared to *U2AF1*^S34F^-NTG (IC_50_: 0.060 µM vs 0.14 µM, p<0.0054) (Figure 5B and Supplementary Figure S4D). Consistent results were obtained in hematopoietic progenitor cells from doxycycline-inducible *U2AF1*^WT^ and *U2AF1*^S34F^ transgenic mice that constitutively express Cas9 (MG-Cas9 mouse cells). *U2AF1*^S34F^-NTG cells were 15-fold more sensitive to CHK1 inhibition than *U2AF1*^WT^-NTG cells (IC_50_ ≈ 0.67 µM vs. 0.10 µM, p<0.0001). *Runx1* depletion made the *U2AF1*^S34F^ cells more resistant, raising the IC_50_ of *U2AF1*^S34F^-sgRunx1 cells 3.3-fold (IC_50_ ≈ 0.67 µM vs. 0.22 µM, p<0.0069) (Supplementary Figures S5C,D).

Next, we tested whether *RUNX1* loss alters CHK1-inhibitor response in human *SRSF2*-mutant leukemia cells. To test this, we used two myeloid leukemia cell lines: KO52 with endogenous *SRSF2*^P95H^ and K562 engineered to express *SRSF2*^P95H^. We depleted *RUNX1* with shRNAs, confirmed knockdown by western blot, and measured GR-normalized dose–response to prexasertib or SRA-737.

K562 experiments. Under non-targeting control shRNA, *SRSF2*^P95H^ K562 showed greater prexasertib sensitivity than parental K562 (Supplementary Figure S6A). *RUNX1* knockdown in *SRSF2*^P95H^ K562 decreased prexasertib sensitivity, with a rightward shift in the GR dose–response relative to control shRNA (Supplementary Figure S6B). RUNX1 protein levels were markedly reduced in *SRSF2*^P95H^ mutant K562 cells treated with *RUNX1*-targeting shRNA (Supplementary Figure S6C).

KO52 experiments. *RUNX1* knockdown reduced KO52 sensitivity to SRA-737 relative to control shRNA, with a rightward shift in the GR dose–response (Supplementary Figure S6D). Western blot confirmed *RUNX1* knockdown (Supplementary Figure S6E).

Collectively, our results show that Runx1/RUNX1 loss reduces sensitivity to CHK1 inhibitors across systems: *Srsf2*^P95H^ and *U2AF1*^S34F^ mouse progenitors, *SRSF2*-mutant patient samples with *RUNX1* mutation, and human leukemia lines with *SRSF2* mutation (K562, KO52).

## DISCUSSION

We found that *SRSF2* mutations create a therapeutically targetable vulnerability to inhibition of ATR, CHK1, and WEE1 DNA damage response kinases. Primary AML samples with *SRSF2* mutations were sensitive to ATR, CHK1 and WEE1 inhibitors, except when *RUNX1* mutations were present. The VAF of *SRSF2* mutations correlated with sensitivity of patient samples to CHK1 and WEE1 inhibitors providing further evidence that *SRSF2* mutant cells have select sensitivity to inhibition of these kinases. In contrast, AML cells from healthy donors were largely refractory. These results suggest a therapeutic window for ATR, CHK1, and WEE1 inhibition in treating *SRSF2* mutant cancers. However, primary AML contain multiple co-occurring mutations, leaving open the question whether the *SRSF2* mutations alone are sufficient to drive sensitivity.

To determine whether *SRSF2* mutations alone are sufficient to sensitize leukemia cells to ATR/CHK1/WEE1 inhibition, we modeled the mutations in two complementary systems: (i) isogenic human K562 cell lines and (ii) knock-in mouse bone marrow progenitors. Using CRISPR/Cas9, we introduced *SRSF2*^P95H/L/R^ alleles into the endogenous locus of K562 cells. Because K562 harbors three copies of *SRSF2*[33], we generated clones with one to three mutant alleles and thus could assess allele-dosage effects. Each mutant clone showed a 2–5-fold decrease in GR_50_ for ATR, CHK1, and WEE1 inhibitors relative to controls, with sensitivity increasing with the number of mutant alleles (Figure 4). In parallel, bone marrow progenitors from *Srsf2*^P95H^ and *U2AF1*^S34F^ knock-in mice displayed heightened sensitivity to CHK1 inhibition (Figure 5), confirming that splicing factor mutations alone create a dependency on DDR kinases.

Prior studies show that mutations in splicing factors lead to aberrant R-loop accumulation because of impaired release of RNA polymerase II from promoter-proximal pausing [6, 32, 39, 40]. R-loops consist of an RNA–DNA hybrid and a displaced single-stranded DNA strand (ssDNA). The presence of ssDNA within unresolved R-loops leads to replication stress and activation of the ATR–CHK1 pathway. ATR signals through CHK1 and WEE1 to activate the G2/M checkpoint to maintain genome integrity [6, 41–43].

Similarly to *SRSF2* and *U2AF1* mutations, *SF3B1*^K700E^ has been shown to elevate R-loop burden, chronically activate ATR–CHK1 signaling, and render cells hypersensitive to CHK1 inhibition [40, 44, 45]. As a result, splicing-factor–mutant cells become dependent on ATR–CHK1 signaling for survival—a dependency corroborated by our data.

*RUNX1* loss confers resistance to CHK1 inhibition in splicing-factor–mutant cells. A subset of *SRSF2-*mutant AML samples that were resistant to CHK1 and WEE1 inhibition had co-occurring *RUNX1* mutations (Figure 2). Together with reports that *RUNX1* mutations are associated with resistance to DNA-damaging chemotherapy in AML [46, 47] these observations led us to test whether *RUNX1* loss reduces sensitivity to CHK1 inhibitors in splicing-factor–mutated hematopoietic progenitors.

To test this, we knocked down *Runx1* in multiple models—c-Kit⁺ bone marrow progenitors from *Srsf2*^P95H^ and *U2AF1*^S34F^ mice, K562 with *SRSF2*^P95H^ and KO52 leukemia cells that naturally harbor *SRSF2*^P95H^—and measured CHK1 inhibitor sensitivity. *Runx1* depletion raised prexasertib IC_50_ four- to six-fold in all models. Thus, intact *RUNX1* appears to be required for the ATR–CHK1 pathway vulnerability created by splicing factor mutations.

We created a panel of 11 isogenic K562 cell line clones that stably express recurrent missense alleles *SRSF2*^P95H/L/R^ from the endogenous locus. We used homology-directed CRISPR/Cas9 to introduce the mutant alleles into the endogenous *SRSF2* locus and confirmed their presence at the DNA level using next-generation sequencing. We also performed deep RNA-seq of the edited clones to characterize splicing alterations caused by the mutant *SRSF2* alleles.

Isogenic K562 clones with *SRSF2*^P95H/L/R^ mutations recapitulate splicing alterations in patients. Prior research has shown that the wild-type *SRSF2* protein binds CCNG and GGNG exonic-splicing-enhancer (ESE) motifs with similar affinity [48]. In contrast, mutant SRSF2 binds CCNG-rich ESE motifs more tightly and GGNG motifs more weakly, driving genome-wide inclusion of CCNG-containing cassette exons and exclusion of GGNG exons in both experimental models and primary MDS/AML samples [28, 34, 49–51]. Consistent with these reports, our isogenic clones have increased inclusion of exons with CCNG and reduced the inclusion of exons with GGNG motifs. Furthermore, we observed higher inclusion of the *EZH2* exon 9b in *SRSF2* mutant clones compared to a wild-type control, which aligns with previous reports [28, 35, 52]. *EZH2* exon 9b is a “poison exon” that contains a conserved premature stop codon that induces nonsense-mediated decay, thus leading to the reduction in EZH2 levels. Collectively these results confirm that our isogenic K562 clones with *SRSF2*^P95H/L/R^ mutations recapitulate known splicing alterations described in MDS/AML patients. The isogenic clones created as part of this study enable direct observation of the effects of *SRSF2* mutations without the confounding effects of other genetic variation.

Our findings indicate that recurrent *SRSF2* mutations in MDS/AML create a selective vulnerability to ATR–CHK1 pathway inhibition. Across patient BM-MNCs and isogenic models, *SRSF2* mutations sensitized cells to ATR, CHK1, and WEE1 inhibitors; *U2AF1* mutations similarly increased sensitivity to CHK1 inhibition, extending the therapeutic scope. These observations align with prior mechanistic work showing that mutant *SRSF2* increases R-loop formation and chronically activates ATR signaling. In contrast, *RUNX1* co-mutations in patients, and *Runx1* loss in splicing-factor– mutant progenitors, reduced response to CHK1/WEE1, nominating *RUNX1* mutations as resistance biomarkers.

In conclusion, mutations in *SRSF2*, *U2AF1*, and *RUNX1* represent candidate, clinically actionable biomarkers to guide stratification for ATR/CHK1/WEE1 pathway inhibitors. These data support clinical evaluation of CHK1 and WEE1 inhibitors, in addition to ATR inhibitors, in splicing-factor– mutant myeloid malignancies, which generally respond poorly to standard chemotherapy.

## ACKNOWLEDGEMENTS

We thank the Helsinki Biobank for the control samples and the Finnish Hematology Registry and Clinical Biobank (FHRB) for providing samples. This work was supported by the Research Council of Finland (grant 334273), the Sigrid Jusélius Foundation, the Magnus Ehrnrooth Foundation, the European Association for Cancer Research (S.E.), the Leukemia and Lymphoma Society (T.G. and M.W.), and NIH P50CA171963 (T.G. and M.W.). M.W. is supported by the Edward P. Evans Foundation and the Lottie C. Hardy Charitable Trust.

## AUTHOR CONTRIBUTIONS

S.E. and T.G. conceived the project and provided leadership. S.E., S.R., T.H., C.C.P, A.B., and A.G. performed experiments. S.R. and A.G. created the isogenic K562 cell line clones. S.E. and V.S., T.H. analyzed and interpreted the data. K.P. collected patient samples and clinical data. S.E., V.S., T. H. and T.G. wrote the paper. All authors critically read the paper, provided constructive comments, and agreed to the content.

## Supplementary information

