## Supplementary Information for "Sensitivity to ATR–CHK1 pathway inhibition in AML/MDS is enhanced by *SRSF2* mutations and reduced by RUNX1 loss"

### Supplementary Materials and Methods

**Patient samples**. Bone marrow samples were collected after informed consent from patients with acute myeloid leukemia (AML) using protocols approved by the Institutional Review Board at the Helsinki University Hospital (permit numbers 239/13/03/00/2010, 303/13/03/01/2011, Helsinki University Hospital Ethics Committee) in compliance with the Declaration of Helsinki. Mononuclear cells (MNCs) were isolated by Ficoll density gradient and resuspended in conditioned medium.

***Ex vivo* drug sensitivity testing of patient cells**. Bone marrow mononuclear cells (MNCs) from healthy donors or patients with AML were cultured in RPMI-1640 media supplemented with 10% fetal bovine serum, 2 mM L-glutamine, penicillin, streptomycin, and 12.5% HS-5 cell line conditioned media [1]. The MNCs were incubated on pre-drugged 384-well plates (Corning) containing up to 510 drugs at five different concentrations in tenfold dilution steps and incubated for 72 h at 37 °C with 5% $CO_2$. Cell viability was measured using the CellTiter-Glo (CTG) 2.0 assay (Promega), using a PHERAstar FS plate reader (BMG LABTECH). Five concentrations were tested, prexasertib: 0.0001–1 µM; PF-00477736, adavosertib, ceralasertib: 0.001–10 µM Dimethyl sulfoxide (DMSO, 0.1%) and benzethonium chloride (100 µM) were used as negative and positive controls, respectively. A four-parameter dose-response curve was fitted, and the drug sensitivity score (DSS), a measure based on the area under the dose-response curve that captures both the potency and the depth of the compound effect, was calculated as previously described [1].

**Identification of mutations and chromosomal aberrations in patient samples.** Gene mutations were identified by whole exome sequencing of leukemic bone marrow and matched skin biopsies.

Whole exome libraries were prepared using NimbleGen SeqCap EZ v2.0, NimbleGen SeqCap EZ MedExome (Roche), Agilent SureSelect Target Enrichment System (Agilent), or Nextera Flex Exome (Illumina) kits according to the manufacturer's guidelines. The exome libraries were sequenced using a HiSeq 2500 instrument (Illumina). Somatic mutations were identified as described previously [2]. To identify chromosomal aberrations, we retrieved clinical karyotype data for AML patient samples from the Finnish Hematology Registry and Biobank, the Hospital District of Helsinki.

**Generation of isogenic gene-edited cell lines.** *SRSF2*$^{P95H/L/R}$ mutations were introduced into the endogenous locus by homology-directed CRISPR/Cas9 editing. K562 (CCL-243; ATCC) cells were resuspended in Buffer R (Thermo Fisher Scientific), gently mixed with a preassembled Cas9-guide RNA ribonucleoprotein (Hifi2-Cas9, MCLAB; gRNA, Synthego) and a single-stranded donor oligodeoxynucleotide repair template (Integrated DNA Technologies), and electroporated using the Neon Electroporation System (Thermo Fisher Scientific). Isogenic *SRSF2*-wildtype clones subjected to identical conditions but lacking edits at the targeted site were retained as controls. Sequences of primers, guides, and templates are provided in **Supplementary Table S1**.

K562 cells were cultured in IMDM medium (Gibco) supplemented with 10% fetal bovine serum, 5000 IU Penicillin/Streptomycin, and glutamate. All cell lines used in this study were tested for Mycoplasma and passaged for less than two months after thawing.

**Detection of *SRSF2*$^{P95H/L/R}$ mutations by sequencing.** We validated the presence of the *SRSF2*$^{P95H/L/R}$ alleles in the isogenic K562 clones by Sanger sequencing. Allele copy number was inferred from variant allele frequencies (VAFs) obtained by targeted amplicon next-generation sequencing (NGS). Primer sequences are listed in **Supplementary Table S3.**

**Cell line viability assays**. Cells were seeded in clear flat-bottom 96-well plates (Corning) at a density of 10,000 cells per well in 100 µl of medium. After 24 h at 37°C, 5% $CO_2$ in a humidified incubator, serial drug dilutions were dispensed using a D300e Digital Dispenser (Tecan). Cells were incubated with the indicated drugs for 72 h. Viability was measured with the CellTiter 96® AQueous One Solution (MTS) assay (Promega) on a SpectraMax M5 plate reader (Molecular Devices) at 490 nm (with a 620–650 nm reference) and normalized to DMSO-treated controls.

To compare drug sensitivity across clones with different growth rates, cell viability measurement results were normalized using the GR metrics R package [3, 4]. Dose-response curves were fit to GR normalized viability values using a four-parameter logistic function in GraphPad Prism v9.0 (GraphPad Software) to obtain $GR_{50}$ values.

**Cell viability by live cell imaging**. Sensitivity to prexasertib in a K562 *SRSF2*$^{P95L}$ mutant clone versus the parental K562 control was assessed using the IncuCyte live cell imaging system (Sartorius). Cells were seeded in 96-well plates and treated with varying concentrations of prexasertib. Real-time cell viability was monitored via the IncuCyte system, which captured and analyzed live cell images at predefined intervals. Viability measurements were normalized using Growth Rate (GR) values to compare drug sensitivity accurately between the mutant clone and the parental control.

**RNA isolation and sequencing**. RNA was extracted from K562 cells using RNeasy Plus Mini Kit (Qiagen). Poly(A)-selected, unstranded Illumina libraries were prepared with the NEBNext Ultra II RNA Library Prep Kit (New England Biolabs). 150 million 150 bp paired-end reads per sample were sequenced using the NovaSeq 6000 (Illumina).

**Mutant allele frequencies in RNA.** To determine *SRSF2* mutant allele frequencies, paired-end reads were aligned to the human reference genome (NCBI GRCh38) using the STAR aligner [5] using default parameters. Sorted, indexed BAMs were reviewed in IGV. At *SRSF2*$^{P95H/L/R}$ sites, we recorded alternate and total read counts and calculated RNA VAF as alt/total.

**Processing and alignment of RNA sequence reads for differential splicing analysis**. To remove sequencing adapters and select 150 bp reads, raw fastq files were processed with Fastp version 0.20.0 with default parameters [6]. To generate a comprehensive reference file of known transcripts, gene annotations in the Ensembl 87 [7] and UCSC knownGene databases [8] were merged. The sequence reads were aligned to the hg19 (NCBI GRCh37) reference genome. The first round of read alignment to the reference of known transcripts consisted of RSEM v1.3.1 [9] using Bowtie v1.0.0 [10] with parameters --bowtie-m 100, --bowtie-chunkmbs 500, --calc-ci and --output-genome-bam. Aligned reads with less than 6 bp overhang across splice junctions and a MAPQ score of zero were filtered out. To maximize the total number of aligned reads spanning splice junctions, unaligned reads from the first step of alignment were realigned with Tophat v2.1.1 [11] using parameters –bowtie1 –readmismatches 2 –read-edit-distance 2 –no-mixed –no-discordant –min-anchor-length 6 –splice-mismatches 0 –min-intron-length 10 –max-intron-length 1000000 –min-isoform-fraction 0 –no-novel-juncs –no-novel-indels –raw-juncs. Reads aligned with Tophat were filtered using the same criteria as in the previous alignment step. The alignment files from RSEM and Tophat were merged. The merged alignment was used for differential splicing analysis.

**Differential splicing analysis**. MISO v2.0 [12] splice event annotations were used to detect splicing changes between mutant and wild-type clones. Only those MISO annotated events that are represented in the set of known reference transcripts described above were used for differential splicing analysis. To evaluate candidates for differential splicing analysis, events with a minimum of 20 informative reads that provide evidence for the presence of specific splice isoforms were selected. Percent Spliced In (PSI) was calculated as the percentage of the read count of a splice isoform over the total read counts of the two alternative isoforms. Differentially spliced events were defined as those that exhibited an absolute isoform ratio difference of 10% and had a Bayes factor of 5, computed using the Wagenmakers' Bayesian framework [13].

**Exonic splicing enhancer motif enrichment in cassette exons**. Biopython [14] was used to identify SSNG exonic splicing enhancer motifs across cassette exon events (where S is C or G and N is any nucleotide). The log2 relative enrichment of a motif was defined as the mean count of instances in included cassette exons divided by the mean count of instances in excluded cassette exons. The relative enrichment was visualized using bar plots.

Ribbon plots were used to visualize the relative frequency of CCNG and GGNG motifs across the differentially spliced cassette exons. The log2 enrichment of the counts of these motifs in included cassette exons were compared to those in excluded cassette exons. A 10-nucleotide sliding window was used for the motif representation. The 95% confidence intervals were calculated using the bootstrapping algorithm in the BRGenomics R package. The results were visualized using the ggplot2 R package.

To visualize splicing events, Sashimi plots were generated using MISO with merged alignment files as input.

**Mouse models**: All experiments were conducted in accordance with institutional guidelines for the care and use of laboratory animals and were approved by the Institutional Animal Care and Use Committee at Washington University in St. Louis. The generation of *Srsf2*$^{P95H/+}$ conditional knock-in mice with *Mx1-Cre* was described by Kim et al. [15]. The humanized *U2AF1*$^{S34F/+}$ (MGS34F) conditional knock-in mice were generated by Fei et al. [16]. Both the *Srsf2*$^{P95H/+}$ and *U2AF1*$^{S34F/+}$ models carried an *Mx1-Cre* transgene enabling the inducible expression of Cre-recombinase, which was induced by intraperitoneal injections of polyinosine-polycytosine (pIpC) at a dosage of 12 µg/g, administered every other day for three days, as previously described[17]. The generation and genotyping of doxycycline-inducible *U2AF1*$^{S34F}$ and *U2AF1*$^{WT}$ transgenic mice were performed as described previously by Shirai et al. [18]. To induce exogenous transgene expression, doxycycline was administered at a concentration of 625 ppm in chow for 5 days, following the protocol outlined by Shirai et al.

**CRISPR/Cas9 mediated *Runx1* knockout**: Synthetic single-guide RNAs (sgRNAs) targeting mouse *Runx1* were selected in the UCSC Genome Browser (mm10, SpCas9 NGG) to maximize predicted on-target activity and minimize off-targets. The *Runx1* sgRNA (5′-TACCTGGTTCTTCATGGCCG-3′) and a non-targeting (NTG) control sgRNA (5′-GCTTTCACGGAGGTTCGACG-3′) were synthesized. The ribonucleoprotein (RNP) complex was formed using Alt-R™ S.p. Cas9-GFP V3 (IDT 10008100) and chemically modified sgRNA in a molar ratio of 1:2.5. The RNP complexes were then nucleofected to the mouse BM cells using 500,000 cells per nucleofection, with a voltage of 1700 V, width 20 ms and pulse 1 in Neon electroporation system (Thermo Fisher USA). The nucleofected cells were immediately placed in pre-warmed transplant medium followed by a 24 h incubation at 37 °C, 5% $CO_2$. GFP-positive cells were sorted using flow cytometry. Approximately 50,000 cells were set aside to measure the editing efficiency using PCR amplicon-based deep sequencing, and the libraries were sequenced using the Illumina MiSeq platform. Data were analyzed using CRISPResso2[19].

**Lentivirus preparation and transduction**: HEK293T cells cultured in DMEM containing 10% fetal bovine serum, 10 mM glutamine, and penicillin (100 IU)/streptomycin (100 µg/ml) were used to produce lentivirus particles. Briefly, HEK293T cells were transfected with 3$^{rd}$ generation lentiviral packaging plasmids including pMDLg (Addgene#12251), pRSV-Rev (Addgene#12253), and pMD.2G (Addgene#12259) using polyethylenimine (Polysciences Inc.) for 12 h, and then the medium was changed with fresh DMEM. Supernatant was collected at 48 h and 72 h, followed by the filtration through a 0.45µm filter. The virus particles were concentrated using Lenti-X™ Concentrator

(Takara Bio, Japan, Cat. 631231). BM cells were transduced with the virus MOI 20, by spinfection at 1200 ×g for 2 h at 30 °C along with polybrene (5ug/mL). Subsequently, 24 h post-transduction, RFP-positive cells were sorted using flow cytometry, and approximately 50,000 cells were kept aside for PCR amplicon-based sequencing, and the libraries were sequenced using the Illumina MiSeq platform and data analyzed using CRISPResso2.

**Immunoblots:** Cells were suspended in Radio Immunoprecipitation Assay (RIPA) buffer, which included 1% Nonidet P40, 0.5% sodium deoxycholate, 0.1% SDS, and protease inhibitors (1 mM Pefabloc, 1 ng/µL aprotinin/leupeptin), along with 10 mM β-glycerophosphate and 1 mM N-ethylmaleimide. Protein concentrations were assessed and equalized using the Pierce BCA Protein Assay Kit (Thermo Fisher Scientific, 23227). Samples were then combined with 4X NuPAGE™ LDS Sample Buffer (Thermo Fisher Scientific; NP0008) containing 50 mM beta-mercaptoethanol (Sigma, M7522) in a 1:4 ratio. The mixture was heated at 95°C for 10 minutes, followed by loading onto polyacrylamide gels and running electrophoresis at 100 V for 90 minutes. Subsequently, proteins were transferred to PVDF membranes for 1.5 h at 250 mA.

The membranes were blocked for 1 h at room temperature using Tris-buffered saline with 0.05% Tween-20 (TBS-T) and 5% milk powder. After blocking, primary antibodies [AML1 (D4A6) Rabbit mAb #8529, Cell Signaling, Monoclonal Anti-β-Actin antibody, Sigma # A5441-2ML] were applied to the membranes and incubated overnight at 4°C. Afterward, the membranes underwent three washes in TBS-T and were treated with secondary antibodies conjugated to horseradish peroxidase for 1 h at room temperature. Following three additional washes, the enhanced chemiluminescence (ECL; Bio-Rad 1705061) substrate was added, and the signals were visualized using the ChemiDoc imaging system (Bio-Rad) with ImageLab v6.0.1 software.

**Statistics**: All data are presented as mean ± standard deviation (SD). Unless otherwise specified, all statistical analyses and graph plotting were carried out using Prism 10 software (GraphPad Software). A p-value of less than 0.05 was considered statistically significant.

### Supplementary Figures

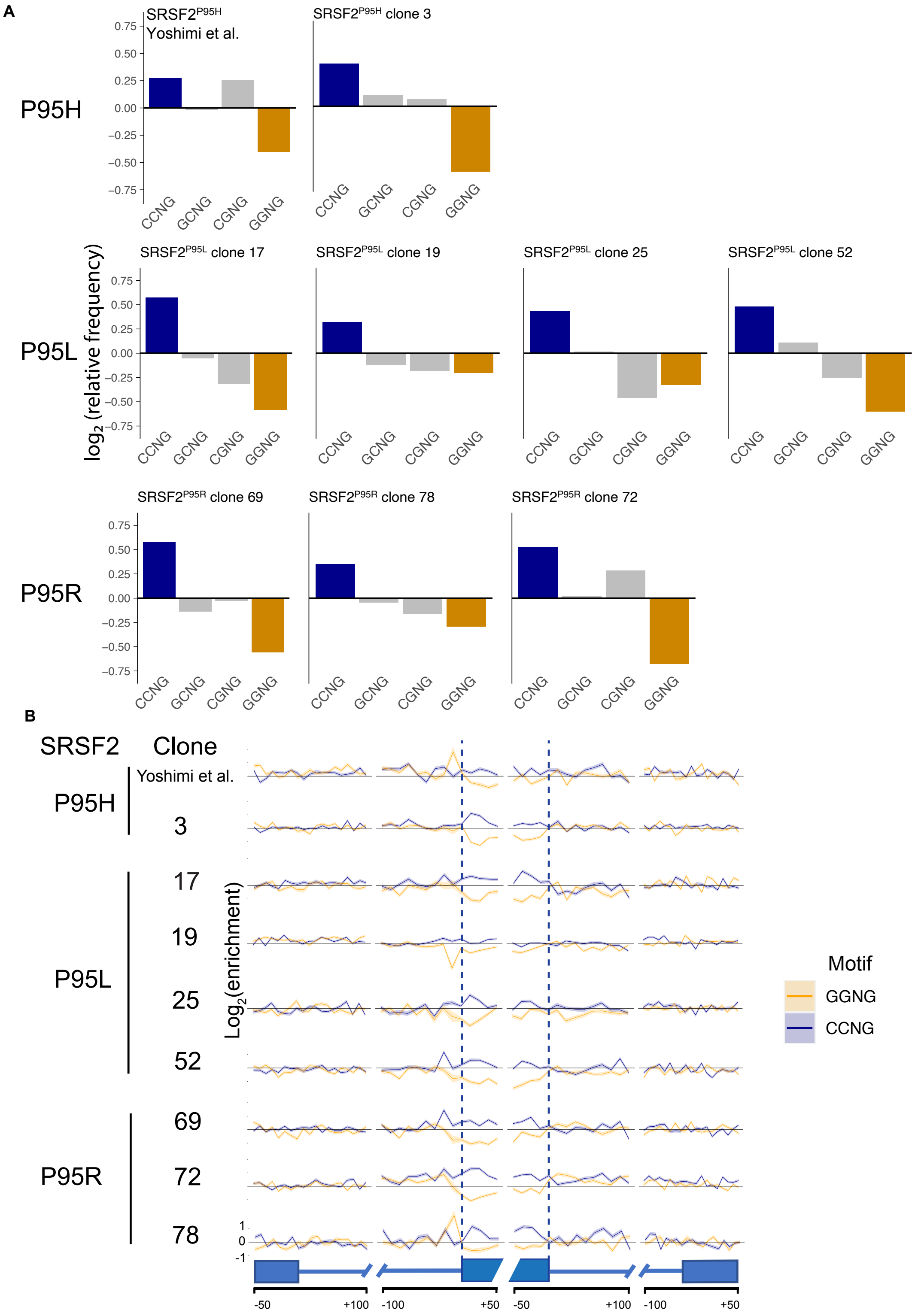

**Supplementary Figure S1. Altered exonic splicing enhancer preference in isogenic K562 clones with *SRSF2*$^{P95H/L/R}$ mutations**. (**A**) Mean enrichment of variants of the SSNG exonic splicing enhancer motif within cassette exons in promoted vs repressed cassette exons in isogenic K562 cells with *SRSF2*$^{P95H/L/R}$ mutations compared to controls. The mean enrichment for a given motif was defined as the mean of the count of instances in all promoted exons divided by the count of instances in all repressed exons. (**B**) Relative frequency of CCNG and GGNG motifs in cassette exons promoted versus repressed by *SRSF2* mutations in isogenic K562 clones. The Y-axis indicates relative motif frequency averaged over promoted vs repressed cassette exons. The middle panels depict the junctions at either end of the cassette exon (diagonally shaded boxes) flanked by immediate junctions of the surrounding exons (blue boxes). Blue lines show adjacent intronic regions. The X-axis is the relative nucleotide position to the respective splice site junction. The shaded areas of the ribbon plots represent >=95% confidence limit obtained through bootstrapping.

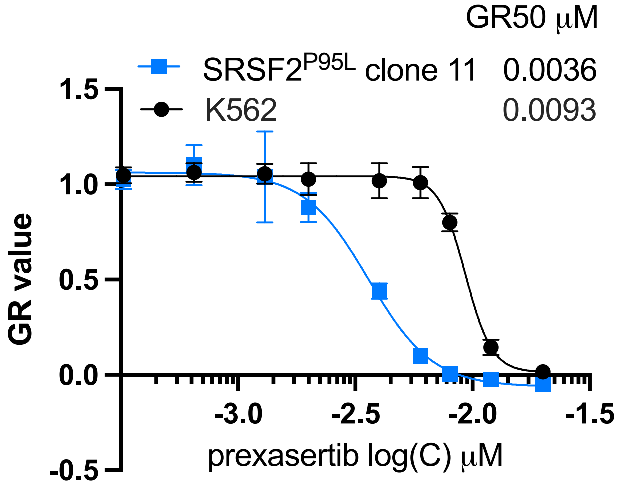


**Supplementary Figure S2: Sensitivity to CHK1 inhibitor in isogenic K562 clone with *SRSF2*$^{P95L}$ mutation measured by live cell imaging.** Prexasertib dose-response of *SRSF2*$^{P95L}$ clone and matched wild-type control. Cell viability was measured by counting cells using an Incucyte Live Cell analysis system after 96 h incubation.

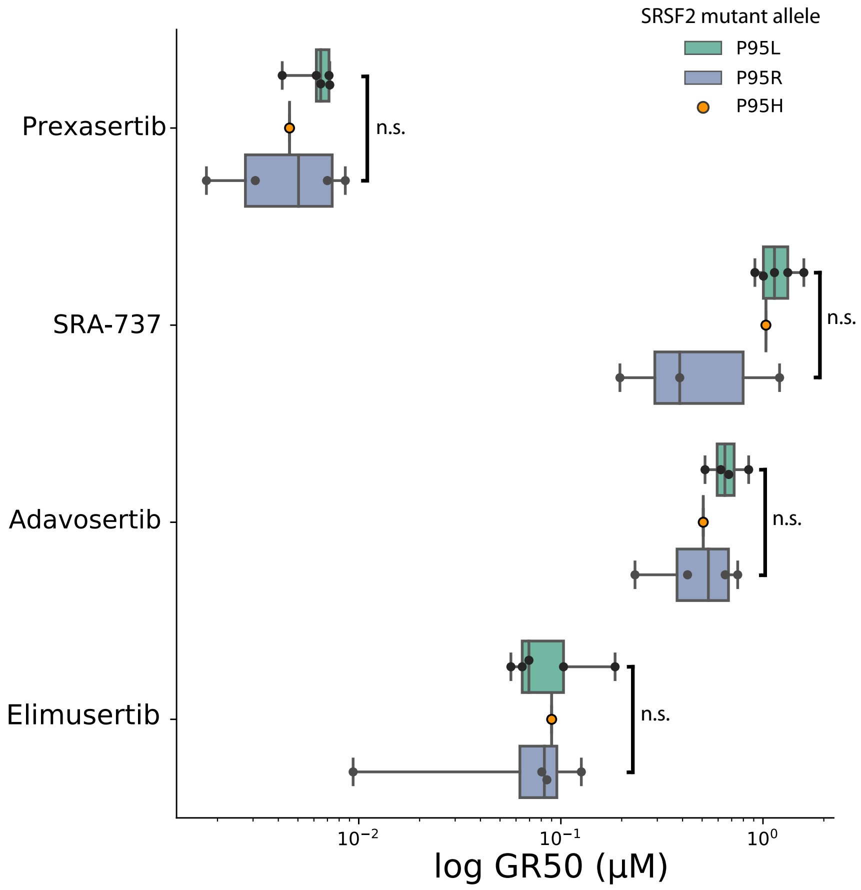


**Supplementary Figure S3: Comparison of drug sensitivity among K562 clones harboring different *SRSF2* mutant alleles.** Sensitivity to ATR, CHK1, and WEE1 inhibitors in a panel of 10 isogenic K562 clones engineered to harbor the *SRSF2*$^{\mathrm{P95H/L/R}}$ mutant alleles and 4 clones with wildtype *SRSF2*. GR normalized half-maximal inhibition to ATR, CHK1, and WEE1 inhibitors across these clones is displayed (x-axis). Clones are grouped according to the *SRSF2* allele they carry. Statistically non-significant (n.s.) by t-test.

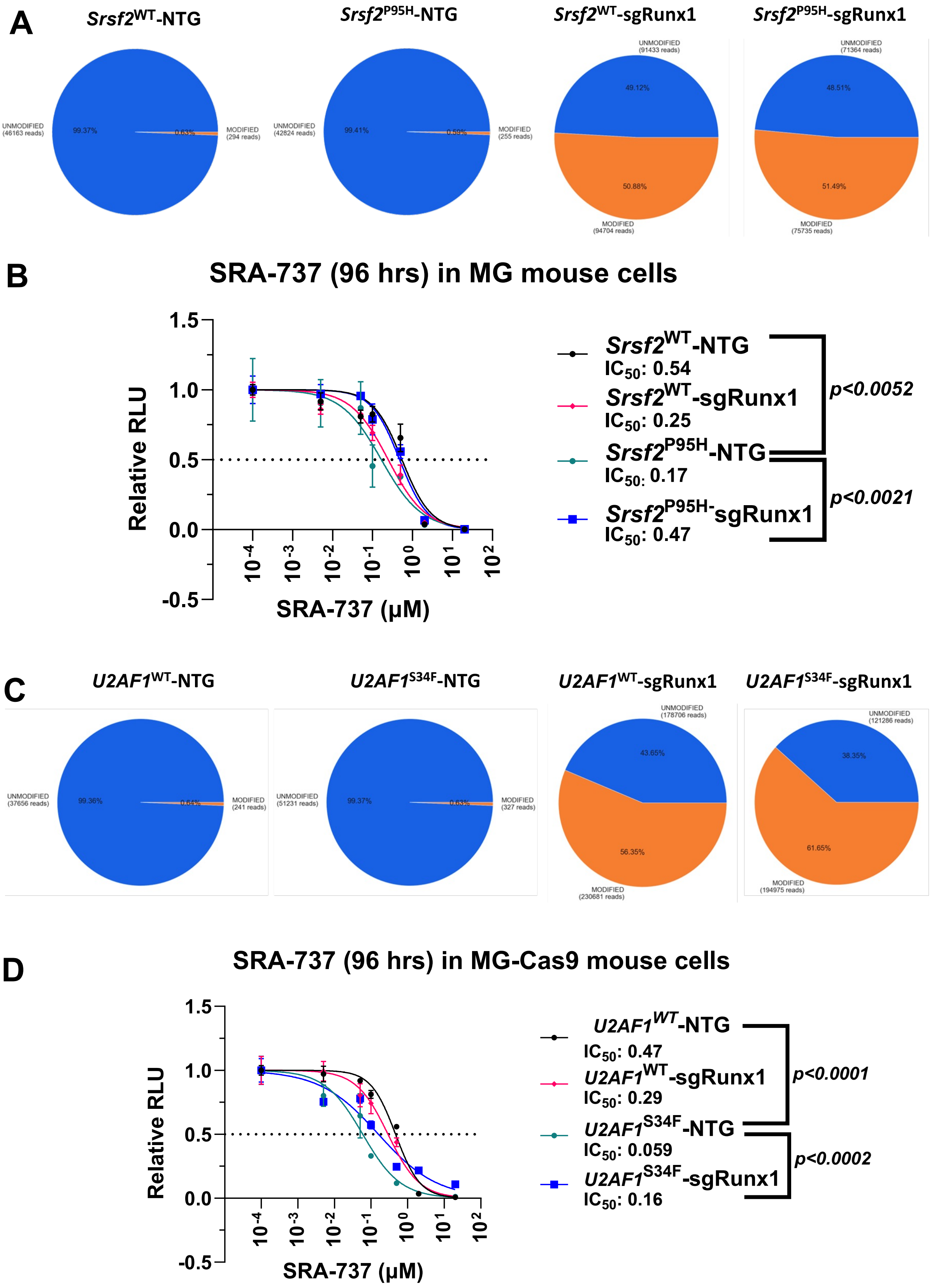

A
Srsf2WT-NTG
Srsf2P95H-NTG
Srsf2WT-sgRunx1
Srsf2P95H-sgRunx1
UNMODIFIED (46163 reads)
99.37%
0.63%
MODIFIED (294 reads)
UNMODIFIED (42824 reads)
99.41%
0.59%
MODIFIED (255 reads)
UNMODIFIED (91433 reads)
49.12%
50.88%
MODIFIED (94704 reads)
UNMODIFIED (71364 reads)
48.51%
51.49%
MODIFIED (75735 reads)
B
SRA-737 (96 hrs) in MG mouse cells
Relative RLU
SRA-737 (μM)
Srsf2WT-NTG IC50: 0.54
Srsf2WT-sgRunx1 IC50: 0.25
Srsf2P95H-NTG IC50: 0.17
Srsf2P95H-sgRunx1 IC50: 0.47
p<0.0052
p<0.0021
C
U2AF1WT-NTG
U2AF1S34F-NTG
U2AF1WT-sgRunx1
U2AF1S34F-sgRunx1
UNMODIFIED (37656 reads)
99.36%
0.64%
MODIFIED (241 reads)
UNMODIFIED (51231 reads)
99.37%
0.63%
MODIFIED (327 reads)
UNMODIFIED (178706 reads)
43.65%
56.35%
MODIFIED (230681 reads)
UNMODIFIED (121286 reads)
38.35%
61.65%
MODIFIED (194975 reads)
D
SRA-737 (96 hrs) in MG-Cas9 mouse cells
Relative RLU
SRA-737 (μM)
U2AF1WT-NTG IC50: 0.47
U2AF1WT-sgRunx1 IC50: 0.29
U2AF1S34F-NTG IC50: 0.059
U2AF1S34F-sgRunx1 IC50: 0.16
p<0.0001
p<0.0002

**Supplementary Figure S4. *Srsf2* mutant, *Runx1* depleted mouse cells exhibit reduced sensitivity to Chk1 inhibition.** (**A, C**) sgRunx1 editing QC**.** Editing efficiency was assessed in bulk c-Kit⁺ bone-marrow cells by PCR-amplicon NGS. Libraries (270 bp) were sequenced on an Illumina MiSeq and analyzed with CRISPResso2 (±5 bp window). Pie charts display unmodified (blue) and modified (orange) reads; read depth and percentage are annotated on each chart. **(B)** CHK1 inhibitor SRA-737 — MG mice**.** Primary c-Kit⁺ bone-marrow cells from doxycycline-induced MG mice (*Srsf2*$^{WT}$-NTG, *Srsf2*$^{WT}$-sgRunx1, *Srsf2*$^{P95H}$-NTG, *Srsf2*$^{P95H}$-sgRunx1) were treated for 96 h with SRA-737 (0.005–20 µM, nine concentrations). The *Srsf2*$^{P95H}$ mutation lowered the $IC_{50}$, indicating increased sensitivity, whereas additional *Runx1* knock-down raised the $IC_{50}$ only in *Srsf2*$^{P95H}$ cells, restoring resistance; *Srsf2*$^{WT}$ cells were unaffected. Viability was measured with CellTiter-Glo 2.0. Points represent mean ± SD of three technical replicates. **(D)** CHK1 inhibitor SRA-737 in MG-Cas9 mouse cells. C-Kit⁺ bone-marrow cells from MG-Cas9 mice (constitutively expressing Cas9) with the *U2AF1* genotypes were treated under the identical conditions described in (B). The response pattern was unchanged: *U2AF1*$^{S34F}$ lowered the $IC_{50}$, and *Runx1* knock-down restored resistance only in the mutant background, while *U2AF1*$^{WT}$ cells remained unaffected. Data are displayed as in (B); pair-wise $IC_{50}$ differences were evaluated by extra-sum-of-squares F-test.

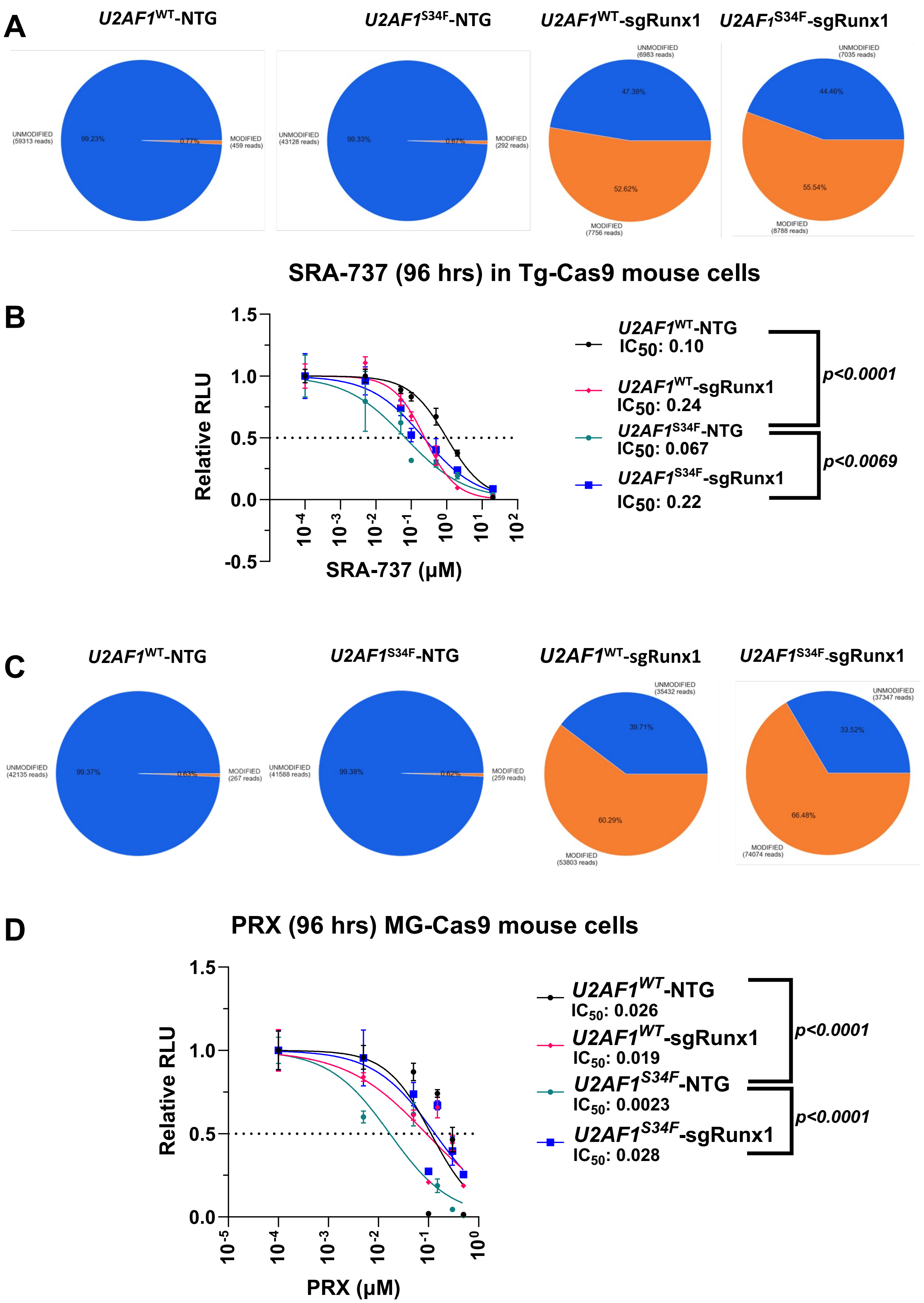
A
U2AF1WT-NTG
U2AF1S34F-NTG
U2AF1WT-sgRunx1
U2AF1S34F-sgRunx1
UNMODIFIED (59313 reads)
99.23%
0.77%
MODIFIED (459 reads)
UNMODIFIED (43128 reads)
99.33%
0.67%
MODIFIED (292 reads)
UNMODIFIED (6983 reads)
47.38%
52.62%
MODIFIED (7756 reads)
UNMODIFIED (7035 reads)
44.46%
55.54%
MODIFIED (8788 reads)
B
SRA-737 (96 hrs) in Tg-Cas9 mouse cells
Relative RLU
SRA-737 (μM)
U2AF1WT-NTG
IC50: 0.10
U2AF1WT-sgRunx1
IC50: 0.24
U2AF1S34F-NTG
IC50: 0.067
U2AF1S34F-sgRunx1
IC50: 0.22
p<0.0001
p<0.0069
C
U2AF1WT-NTG
U2AF1S34F-NTG
U2AF1WT-sgRunx1
U2AF1S34F-sgRunx1
UNMODIFIED (42135 reads)
99.37%
0.63%
MODIFIED (267 reads)
UNMODIFIED (41588 reads)
99.38%
0.62%
MODIFIED (259 reads)
UNMODIFIED (35432 reads)
39.71%
60.29%
MODIFIED (53803 reads)
UNMODIFIED (37347 reads)
33.52%
66.48%
MODIFIED (74074 reads)
D
PRX (96 hrs) MG-Cas9 mouse cells
Relative RLU
PRX (μM)
U2AF1WT-NTG
IC50: 0.026
U2AF1WT-sgRunx1
IC50: 0.019
U2AF1S34F-NTG
IC50: 0.0023
U2AF1S34F-sgRunx1
IC50: 0.028
p<0.0001
p<0.0001

**Supplementary Figure S5. *U2AF1*, *Runx1* depleted mouse cells exhibit reduced sensitivity to Chk1 inhibition. (A, C) sgRunx1 editing QC.** The editing efficiency of the sgRNA was tested by PCR amplicon based next generation sequencing. The libraries were sequenced using the Illumina MiSeq platform and data analyzed using CRISPResso2. Pie charts display unmodified (blue) and modified (orange) reads; read depth and percentages are shown in each chart. **(B) CHK1 inhibitor SRA-737.** Primary c-Kit$^{+}$ bone-marrow cells from doxycycline-induced *U2AF1*$^{WT}$-NTG, *U2AF1*$^{WT}$-sgRunx1, *U2AF1*$^{S34F}$-NTG, and *U2AF1*$^{S34F}$-sgRunx1 mice were treated for 96 h with SRA-737 (0.005–20 µM, nine concentrations). The *U2AF1*$^{S34F}$ mutation lowered the $IC_{50}$, indicating increased sensitivity, whereas additional Runx1 knock-down raised the $IC_{50}$ only in *U2AF1*$^{S34F}$ cells, restoring resistance; *U2AF1*$^{WT}$ cells were unaffected. Viability was measured with CellTiter-Glo 2.0. Points show mean ± SD of three technical replicates. Pairwise $IC_{50}$ differences were evaluated with XX test. **(D) Dual CHK1/2 inhibitor prexasertib.** Using the same induction protocol and cell panel, cultures were exposed for 96 h to prexasertib (0.005–20 µM, nine concentrations). Results paralleled those with SRA-737: *U2AF1*$^{S34F}$ increased drug sensitivity (lower $IC_{50}$), and Runx1 knock-down restored resistance exclusively in the mutant background, with no effect in *U2AF1*$^{WT}$ cells. Data presentation as in (B).

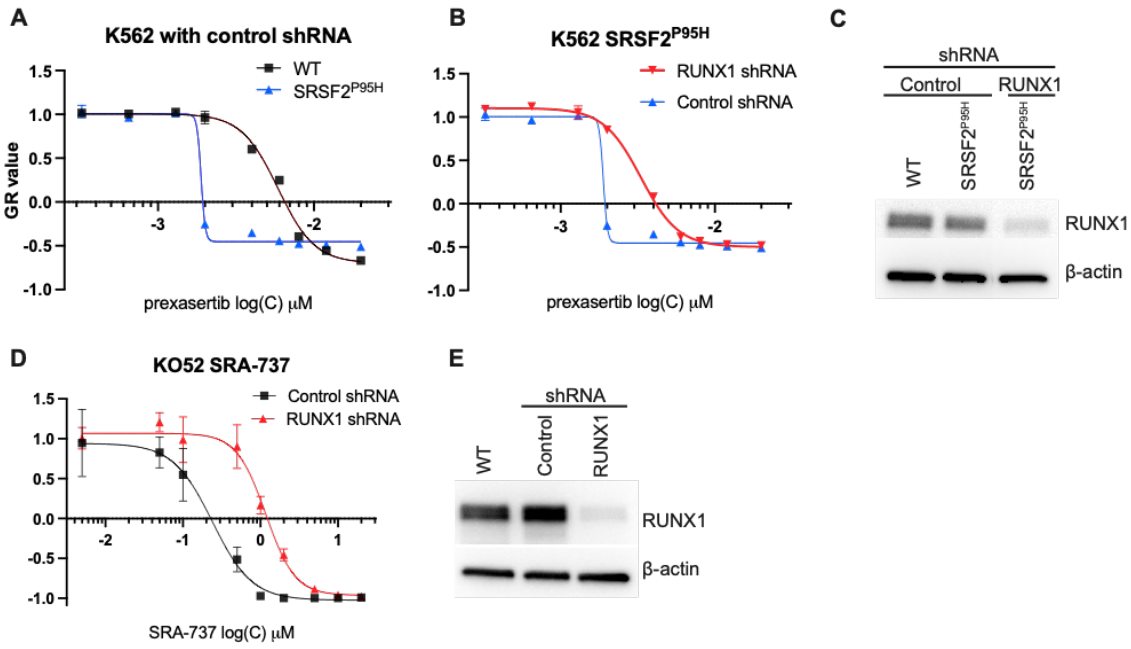


**Supplementary Figure S6. *RUNX1* knockdown induces resistance to CHK1 inhibition in *SRSF2* mutant leukemia cell lines. (A)** Dose-response curves for prexasertib in parental and K562 *SRSF2*$^{P95H}$ K562, both with non-targeting shRNA. Viability at 72 h was measured with CellTiter-Glo and converted to growth-rate–normalized values (GR), which are plotted vs log10 prexasertib concentration. **(B)** Prexasertib sensitivity of K562 cells expressing *SRSF2*$^{P95H}$ with *RUNX1*-targeting or control shRNA. **(C)** Western blot analysis of RUNX1 and β-actin protein levels in K562 clones treated with control or *RUNX1* shRNA in wild-type (WT) and *SRSF2*$^{P95H}$ backgrounds. β-actin is the loading control. **(D)** SRA-737 sensitivity of KO52 cells treated with control shRNA and *RUNX1* shRNA. **(E)** Western blot of RUNX1 and β-actin protein levels in KO52 cells treated with control or *RUNX1*-targeting shRNA.
